# Sampling principles for biodiversity study

**DOI:** 10.1101/000364

**Authors:** Xubin Pan

## Abstract

Sampling is a fundamental tool in ecology and critical for biodiversity measurement. However, basic principles of biodiversity sampling have been overlooked for many years. In this paper, I proposed and explored five principles of sampling for a specific area and biodiversity study. The first principle of sampling, *species increasing with area*, is that the number of species increases with the area. The second principle of sampling, *individuals increasing with area*, is that the number of individuals increases with the area. The third principle of sampling, *sum of species number*, is that the sum of species number in one area and species number in another area is no less than the total species number in the two areas. The fourth principle of sampling, *individual complement*, is that the sum of the mathematical expectation of individual number of one or several species in the area *a* and that of the same one or several species in the area *A-a* is the total individual number *N* of the same one or several species in the total area *A*. The fifth principle of sampling, *species-area theory*, is that the sum of the mathematical expectation of number of species in the area *a* and that of number of species lost if area *A-a* is cleared is the total species number *M* in the total area *A*.

## Introduction

Sampling is a fundamental tool in ecology. Because the complete investigation of one ecological question maybe involves a large area and long time, it will cost too much time and money, and leads to infeasible in practice. Thus, sampling is unavoidable in the ecology study. However, due to the complexity of ecosystem, such as interactions among organisms and environment and their large spatial-temporal heterogeneity, how to sampling should be very carefully, which will decide how to analyze the data from the sampling and the credibility of the results. Thus, solid sampling theory is very important for ecology, although it didn’t get too much concern recently. As the core of the ecology, the spatial relationship between the species and the area can be a good start for rethink the sampling theory.

Individual–area relationship (IAR), species-area relationship (SAR) and endemics-area relationship (EAR) are important concepts in biodiversity conservation and habitat preservation (WCMC, 1992; Kinzig and Harte, 2000; Connor et al., 2000; Green and Ostling, 2003; Chris et al., 2004; World Resources Institute, 2005). However, there is still much debating over the estimation of extinction rate based on SAR, which tends to make overestimation when compared with observed extinction (Pimm and Askins, 1995; Rosenzweig, 1995; Harte and Kinzig, 1997; He and Hubbell, 2011; Pan, 2013). One explanation is that such estimation includes certain species that are “committed to extinction” instead of going extinct right away after habitat clearing (Heywood, 1994; Tilman, 1994; Mace et al., 2003). Recently, He and Hubbell (2011) suggested that the reason was the difference of between sampling areas based SAR and EAR, because a sample area of a species for extinction is often larger than a sample area of the same species for existence. Among these controversies, the core problem is the sampling of biodiversity measurement, the basic principles of which have been overlooked for many years.

However, much work has been focused on statistical or mathematical calculation based on SAR, rather than the biological and ecological implication, especially the basic principles for biodiversity sampling (Turner and Tjørve, 2005). Are there common laws for sampling in biodiversity measurement? Here, we proposed five sampling principles for biodiversity study in a specific area and the last two were proved, focusing on the change of species number and individuals for one or several species with a changing area. This analysis will be helpful for the establishment of theory platform for biodiversity and other ecological discussion.

## Theoretical Frame

Biodiversity not just study the number of species, but also the amount of individuals. For the sampling problems of biodiversity, is the relationship between the area and the number of species and amount of individuals. Usually, it is thought the relationship between the number of species and the area is saturation curve, and the relationship between the amount of individuals and the area is increasing curve. The shape of this curve is influenced by the spatial distribution and sampling collection/statistic.

## Sampling principles for biodiversity measurement

The first principle of sampling, species increasing with area, is that the number of species increases with the area. Assume the number of species is *m* in the area *a*, when the sampling area increases from *a* to *a*′, the number of species in area *a*′ is *m*′, and *m*′ ≥ *m*. If no new species emerges in the area *a*′ − *a*, *m*′ = *m*.

The second principle of sampling, individuals increasing with area, is that the number of individuals increases with the area. Assume the number of individuals is *n* in the area *a*, when the sampling area increases from *a* to *a*′, the number of individuals in area *a*′ is *m*′, and *n*′ ≥ *n*. If there no new individuals emerges in the area *a*′ − *a*, *n*′ = *n*.

The third principle of sampling, sum of species number, is that the sum of species number in one area and species number in another area is no less than the total species number in the two areas. Assume the number of species is *m*_1_ in the area *a*_1_, the number of species is *m*_2_ in the area *a*_2_, the number of species in area *a*_1_ + *a*_2_ is *m*′, and m′ ≤ m_1_ + m_2_. If there is no overlapping species between areas *a*_1_ and *a*_2_, *m*′ = *m*_1_ + *m*_2_.

The fourth principle of sampling, individual complement, is that the sum of the mathematical expectation of individual number of one or several species in area *a* and that of individual number of the same one or several species in area *A-a* is the total individual number *N* of the same one or several species in the total area *A*.

The fifth principle of sampling, species-area theory, is that the sum of the mathematical expectation of number of species in area *a* and that of number of species lost if area *A-a* is cleared is the total species number *M* in the total area *A*.

## Mathematical proof for the fourth and fifth principles of biodiversity sampling

While the first, second and third principles of biodiversity sampling are straightforward and easy to understand, a complementary method in a specific area is used to verify the fourth and fifth principles.

The individual-area relationship (IAR) *I_a_* in area *a* and *I_A-a_* in area *A-a*, for one sampling, if the sampling area is *a* and the number of individuals in this area is *n*, there is always one-one corresponding area of *N-n* individuals of the species in area *A-a* (*A* is the total area, and *N* is the total individuals for one or several species in area *A*).

The mathematical expectation of individual-area relationship in area *a* is

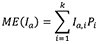

where *ME*(*I_a_*) are the expected individuals for the individuals-area relationship in area *a* in the *i*th sampling, *k* is the total samples, and the *P_i_* is the possibility of the corresponding *i*th sampling.

The mathematical expectation of individual-area relationship in area *A-a* is

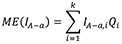

where *ME*(*I_A-a_*) are the expected individuals for the individual-area relationship in area *A-a* in the *i*th sampling, and the *Q_i_* is the possibility of the corresponding *i*th sampling.

For any specific sampling *i*, there is always a one-one corresponding *I_a,i_* + *I_A-a,i_* = *N*, and the *P_i_* = *Q_i_* in this situation.

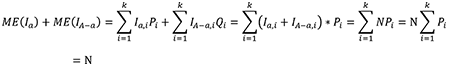

The number of species in area *a* is *m*, then *S_a_ = m*, where species-area relationship (SAR) *S_a_* is the function of the species number with area *a*. The number of species lost in area *A-a* if the area *A-a* is cleared is *M-m*, endemics-area relationship (EAR) *E_A-a_* is the number of species disappearing if area *A-a* is cleared.

The mathematical expectation of Species-area relationship in area *a* is

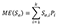

where *S_a,i_* is the species number in area *a* in the *i*th sampling, *k* is the total samples, and the *P_i_* is the possibility of the corresponding *i*th sampling.

The mathematical expectation of Endemics-area relationship in area *A-a* is

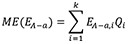

where *E_A-a, i_* is the number of species disappear if area *A-a* is lost in the *i*th sampling, and the *Q_i_* is the possibility of the corresponding *i*th sampling.

For any specific sampling *i*, there is always a one-one corresponding *S_a,i_* + *E_A-a,i_* = *M*, and the *P_i_* = *Q_i_* in this situation.

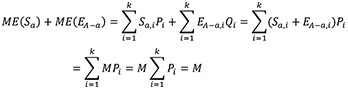

## Acknowledgments

I warmly thank Fengqiao Liu at Arizona State University and Dr. James Rosindell in University of Leeds for the helpful suggestions for the manuscript revision.

**Figure 1.**
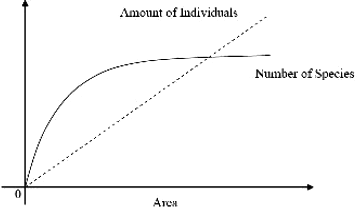
Relationship between the number of species and amount of individuals and the area.

**Figure 2.**
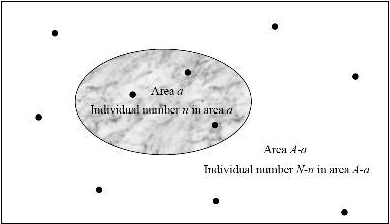
Individual-area relationship *I_a_* in area *a* and *I_A-a_* in area *A-a*.

**Figure 3.**
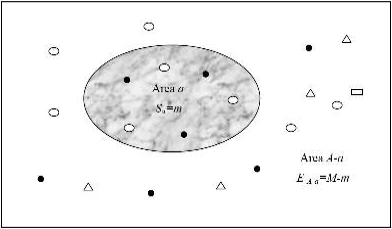
Species-area relationship *S_a_* in area *a* and endemics-area relationship *E_A-a_* in area *A-a*.

